# Pervasive adaptation in *Plasmodium*-interacting proteins in mammals

**DOI:** 10.1101/081216

**Authors:** Emily R. Ebel, Natalie Telis, Sandeep Venkataram, Dmitri A. Petrov, David Enard

## Abstract

The protozoan genus *Plasmodium* causes malaria in dozens of mammal species, including humans, non-human primates, rodents, and bats. In humans, *Plasmodium* infections have caused hundreds of millions of documented deaths, imposing strong selection on certain populations and driving the emergence of several resistance alleles. Over the deep timescale of mammalian evolution, however, little is known about host adaptation to *Plasmodium*. In this work, we expand the collection of known *Plasmodium*-interacting-proteins (PIPs) in mammalian hosts from ~10 to 410, by manually curating thousands of scientific abstracts. We use comparative tests of adaptation to show that PIPs have experienced >3 times more positive selection than similar mammalian proteins, consistent with *Plasmodium* as a major and long-standing selective pressure. PIP adaptation is strongly linked to gene expression in the blood, liver, and lung, all of which are clinically relevant tissues in *Plasmodium* infection. Interestingly, we find that PIPs with immune functions are especially enriched for additional interactions with viruses or bacteria, which together drive a 3.7-fold excess of adaptation. These pleiotropic interactions with unrelated pathogens, along with pressure from other *Plasmodium*-like Apicomplexan parasites, may help explain the PIP adaptation we observe in all clades of the mammalian tree. As a case study, we also show that alpha-spectrin, the major membrane component of mammalian red blood cells, has experienced accelerated adaptation in domains known to interact specifically with *Plasmodium* proteins. Similar interactions with *Plasmodium*-like parasites appear to have driven substantial adaptation in hundreds of host proteins throughout mammalian evolution.

## Introduction

Malaria is one of the world’s most notorious infectious diseases, responsible for billions of illnesses and millions of deaths in the last fifty years alone (WHO, 2015). The malaria genus *Plasmodium* contains five species infecting humans, including *P. falciparum*, and 53 species infecting non-human primates, rodents, and bats (Carlton, Perkins, and Deitsch, 2013). Other blood-borne Apicomplexans, which are frequently confused for *Plasmodium,* also cause malaria-like symptoms in livestock and pets (Escalante and Ayala, 1995; Coatney and Roudabush, 1936; Clark and Jacobson, 1998). The strict genus *Plasmodium* is thought to have experienced a major radiation 55-129 million years ago (Escalante and Ayala, 1995), indicating a long-standing relationship between the malaria parasite and its mammalian hosts.

This ancient relationship raises the possibility that *Plasmodium*, along with similar parasites, have imposed an important and complex selective pressure on mammals. The host-parasite interactions involved in malaria span multiple stages and tissue types, each of which may be subject to selection. Briefly, after the bite of an infected mosquito transmits *Plasmodium* cells into the blood, they migrate to the liver and multiply many times. After several days, parasites emerge from the liver and begin infecting red blood cells (RBCs). The ensuing 48-hour cycles of replication and emergence from RBCs are responsible for anemia, fever, and other characteristic symptoms of malaria. *Plasmodium* parasites are also known to sequester in certain organs, including the brain, lungs, and adipose tissues, which can result in severe complications (Idro et al. 2010; Franke-Fayard et al., 2005; Lovegrove et al., 2008; Aursudkij et al., 1998). Furthermore, parasitic proteins and by-products solicit a complex immune response, including the tagging of parasitized RBCs for removal from circulation by the spleen (Engwerda *et al.*, 2005).

Given these many facets of host-parasite interaction, as well as the substantial morbidity and mortality of malaria, it seems likely that *Plasmodium* has imposed an important selective pressure on its hosts. Over the deep time scale of mammalian evolution, this hypothesis has not yet been tested, but it has been supported over the shorter time scale of human evolution. In African and Southeast Asian populations, several malaria resistance variants appear to have risen in frequency over the last 5,000-10,000 years (Hedrick, 2011; Kwiatkowski, 2005). While some, such as the Duffy null mutation, have approached local fixation (Welch, 1977), many others are prevented from fixing by their deleterious pleiotropic effects. For example, the hemoglobin sickle cell allele offers substantial protection against malaria, but causes fatal anemia in the homozygous form (Aidoo *et al.,* 2002). The fact that balancing selection maintains such a deleterious allele, at up to 15% frequency in some African populations (Piel *et al.,* 2010), suggests that malaria presents a strongly opposing selective force. Indeed, malaria has repeatedly been labeled "one of the strongest selective forces on the human genome" (Hedrick, 2011; Verra *et al*., 2009; Kwiatkowski, 2005), though this statement has never been quantified.

The evolutionary impact of a complex selective pressure is difficult to quantify precisely. One important reason is that the many genes relevant to one phenotype, like malaria resistance, are each likely to be pleiotropically involved with a set of other phenotypes (Wagner and Zhang, 2011). Thus, it is difficult to ascribe evolutionary patterns in a small number of genes with malaria-related functions, such as certain RBC or immune proteins, specifically to the selective pressure imposed by malaria. Limiting evolutionary analysis to certain pathways or genes can also exclude the effects that a complex selective pressure has on other biological systems (see Travisano and Shaw, 2012). Both of these issues could be circumvented, in the case of malaria, by comparing a large set of genes known to interact with *Plasmodium* to a large set of genes that share similar properties, but do not interact with *Plasmodium*. With a sufficient number of genes, this approach should average out various pleiotropic effects, as well as allow for expression in particular tissues to be tested for association with adaptation. A similar approach to gene set enrichment has previously been used to describe polygenic adaptation to pathogens in humans (Daub et al. 2013). More recently, it has been combined with manual curation and carefully chosen controls to identify viruses as a dominant driver of adaptation in mammalian proteins (Enard et al., 2016).

In order to estimate an evolutionary effect specifically attributable to *Plasmodium*, the gene enrichment strategy outlined above necessarily requires a large number of *Plasmodium*-interacting genes. However, only a few dozen such genes had previously been compiled (e.g. Verra et al., 2009), and only ~10 of these are highly conserved across mammals. In this work, we manually examine over 30,000 publications to identify 410 conserved, mammalian *Plasmodium*-interacting proteins, or PIPs. By leveraging extensive collections of functional data from over 9,000 mammalian proteins, we fairly compare rates of evolution between PIPs and control genes, which have been matched to PIPs across a wide range of properties. We find evidence of unusually strong and pervasive positive selection in PIPs, which has likely been driven by interactions with *Plasmodium*-like parasites over millions of years.

## Results

### *Robust identification of 410* Plasmodium*-interacting proteins (PIPs)*

Given the importance of malaria to human populations, research into host-parasite interactions and host health effects has been underway for over a century. We queried the PubMed database for all scientific papers whose abstracts mentioned ‘malaria,’ ‘*falciparum*,’ or ‘*Plasmodium,*’ along with the name of a host gene (Methods I). To focus on mammalian evolution, we limited our search to 9,338 protein-coding genes that are conserved in 24 mammalian species, including humans (Methods II; Enard et al., 2016). Most of these mammalian species belong to one of four orders—primates, rodents, artiodactyls, or carnivores (S1 Fig)—and represent a variety of lifestyles and susceptibilities to malaria.

This search returned 30,788 papers associated with 2,249 of the 9,338 well-conserved genes. However, the vast majority of these results were false positives, due largely to multiple meanings of the short acronyms that identify genes. We manually curated the results to identify 484 papers linking 410 proteins to malaria through any of four types of phenotypic evidence: (1) physical interaction between the mammalian and *Plasmodium* protein; (2) statistical association with malaria susceptibility; (3) knock-out or overexpression studies; and (4) low-throughput studies showing a change in gene expression during malaria infection (Fig 1A; S1 Table). Expression changes were the most common form of evidence (72% of PIPs), but 28% of PIPs were supported by multiple sources of evidence, and 41% by multiple studies. Virtually all of the studies were conducted on five *Plasmodium* species infecting humans or mice (Fig 1A).

**Fig. 1.**
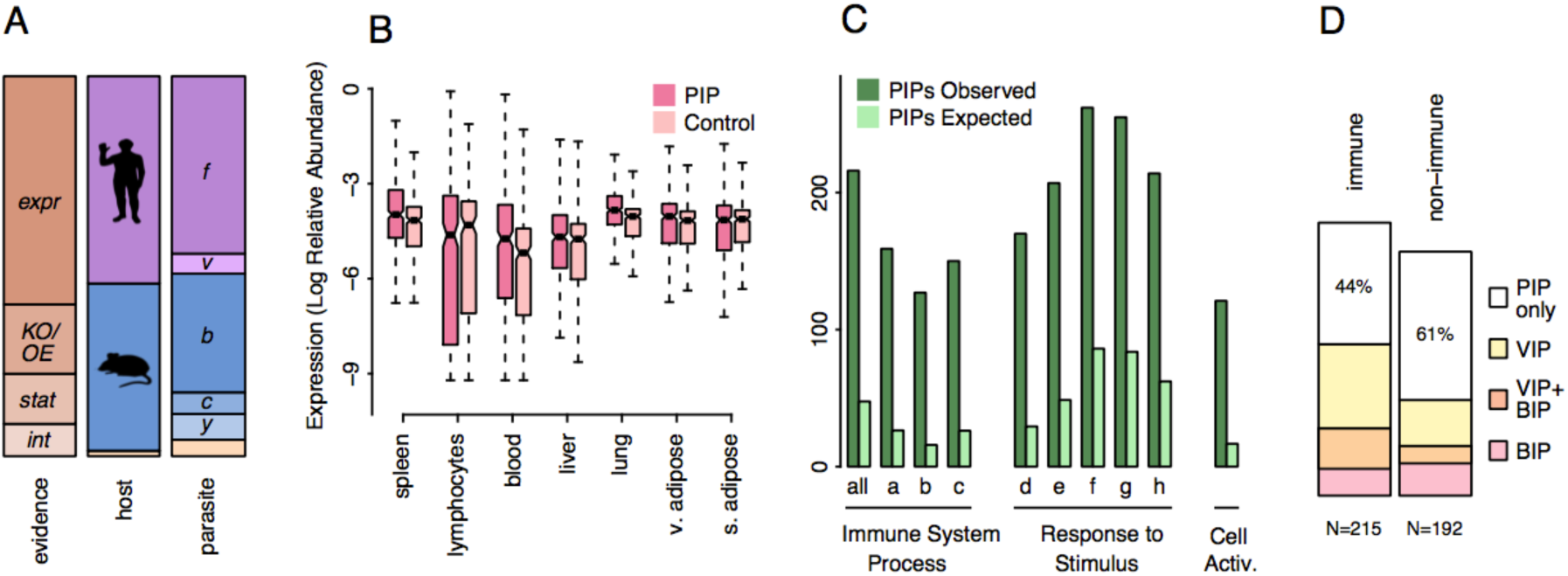
PIP sources, expression patterns, and GO functions. (a) The 410 PIPs are summarized in each stacked bar. Abbreviations: *expr* – expression; *KO/OE* – knockout/overexpression; *stat* – statistical association; *int* - physical interaction; *f – falciparum; v – vivax; b – berghei; c – chabaudi; y – yoelii*. (b) Relative abundance of PIP expression is higher than expected in seven tissues (all p<0.006), compared to control genes matched for total expression. All tissues are shown in S2 Fig. Abbreviations: *v* – visceral; *s* – subcutaneous. (c) Top 10 GO categories most significantly enriched for PIPs (all p<<0.001). Sub-categories a-h are enumerated in S3 Fig. (d) The pathogen pleiotropy of immune PIPs (53% of all PIPs) and non-immune PIPs (47% of all PIPs) is summarized in each stacked bar. 44% of immune PIPs, versus 61% of non-immune PIPs, are not known to interact with any other viral (VIP) or bacterial (BIP) pathogen.

### PIP expression and function support role in malaria

If PIPs are truly a set of malaria-relevant genes, we would expect the pathophysiology of malaria to be reflected in their tissue expression profiles. We tested this hypothesis by examining human gene expression in each of the 53 tissues collected by the GTEx Consortium (2015; Methods III). We first found that, on average, PIPs have 9.5% higher total expression than other genes (p<0.0001; Fig 2A). To fairly evaluate PIP overexpression in each tissue, we designed a matched permutation test that compares PIPs to many, similarly-sized sets of control genes with similar total expression (Methods III). After controlling for total expression in this way, we find seven tissues in which PIPs are significantly overexpressed (Fig 1B, all p<0.006; all tissues are shown in S2 Fig). These include blood and liver, where *Plasmodium* parasites reproduce; spleen and lymphocytes, which participate in the immune response; and the lung and adipose tissues, which are secondary sites of parasite sequestration (Franke-Fayard et al., 2005).

**Fig. 2.**
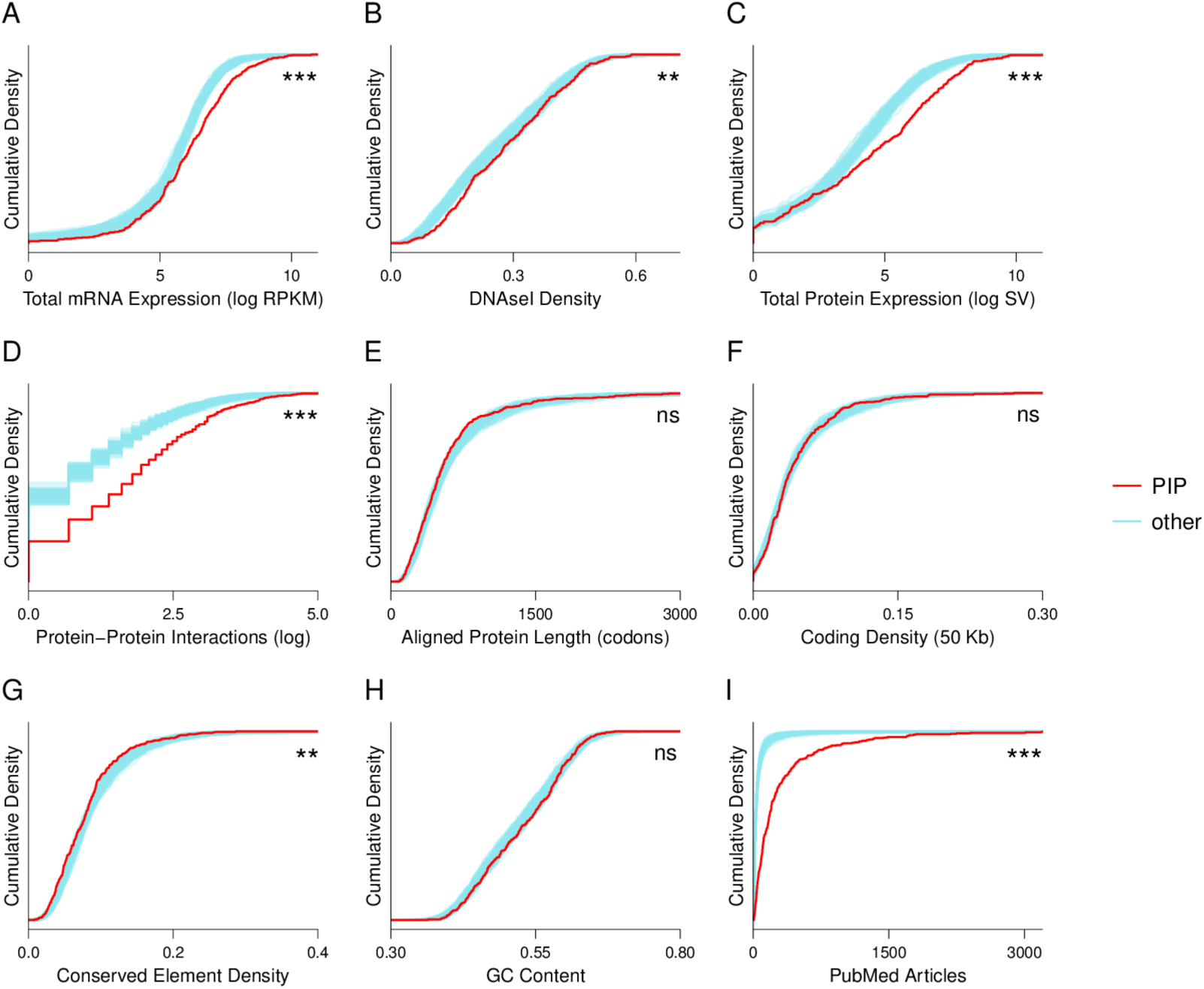
PIPs differ from other conserved genes by a number of functional measures, but meet expectations for most measures of genomic context. Each plot is a cumulative density function, which tracks the proportion of the gene set (y-axis) with values greater than or equal to each value on the x-axis. That is, sets with higher average values will traverse more x-axis space (and appear as ‘lower’ lines) before reaching the maximum density of 100%. Descriptions of data sources are available in Methods V. *** = p<0.0001; ** = p<0.001; ns = p>0.05.

Similarly, we expected PIPs to be enriched for GO functions that reflect malaria pathology. We tested 17,696 GO functional categories (Methods V) for PIP enrichment using Fisher’s Exact Test. After correcting for multiple testing, over 1,000 categories contained significantly more PIPs than expected (S2 Table). These categories are dominated by immune functions, especially for the highest levels of enrichment (Fig 1C). Other functions, including apoptosis, cell-cell signaling, and coagulation, are also highly enriched for PIPs (S2 Table). These results confirm the biological connections between PIPs and malaria, and suggest that immune pathways present a major functional interface between host and parasite.

### Immune and other PIPs are enriched for viral and bacterial interactions

Many immune genes, even outside the adaptive immune system, are known to be activated by signals from multiple pathogens (e.g. Ozinsky et al, 2000; Yamamoto et al., 2013). Such ‘pathogen pleiotropy’ poses an important complication when testing the link between *Plasmodium,* as a single causal pathogen, and adaptation in any gene. To quantify the extent of this pleiotropy for PIPs, we compiled mammalian proteins known to interact with viruses, bacteria, and any Apicomplexan parasites outside the genus *Plasmodium* (Methods IV). For viruses, we obtained a high-quality list of 1,256 manually curated virus-interacting proteins (VIPs) from Enard *et al* (2016). For bacteria and Apicomplexans, we queried the EBI IntAct database (Orchard et al., 2014) for all deposited interactions (see Methods IV). This search returned 1,250 mammalian bacteria-interacting proteins (BIPs; S3 Table), but 0 instances of mammalian interaction with non-*Plasmodium* Apicomplexans. Most Apicomplexan parasites are still poorly studied at the molecular level (Carlton, Perkins, and Deitsch, 2013), but it is likely that parasites in the same phylum as *Plasmodium* have more overlapping interactions with PIPs than do viruses or bacteria.

Overall, we find that 37% of all PIPs also interact with viruses, 22% with bacteria, and 48% with viruses and/or bacteria—many more than expected by chance (all p<0.0001; S6 Fig). As expected, this multi-pathogen overlap is strongest for immune PIPs (Fig 1D). Surprisingly, however, we find that nearly 40% of non-immune PIPs also interact with these unrelated groups of pathogens (Fig 1D). While some of these non-immune, multi-pathogen PIPs could in theory have uncharacterized immune functions, most are known for their involvement in general cellular processes, including metabolism and signal transduction. This suggests that a diverse array of prokaryotic, eukaryotic, and viral pathogens take advantage of a limited number of cellular pathways to infect their hosts. Such pleiotropy has many interesting implications, including the need to carefully isolate any single selective pressure when linking it to protein adaptation.

### PIPs are not like other proteins

We have already shown that PIPs have two unusual properties—high mRNA expression, and excess overlap with other pathogens—that may influence their rate of evolution. We assessed several additional metrics for differences between PIPs and other proteins, in order to fairly evaluate PIP adaptation.

First, we tested three more broad measures of gene function in humans: the density of DNAseI hypersensitive elements; protein expression, as measured by mass spectrometry; and the number of protein-protein interactions (see Methods V). For each of these metrics, PIPs have significantly higher mean values than sets of random controls, indicating that PIPs are more broadly functional in humans (Fig 2, B-D; all p<0.01). We next tested four measures of genomic context, which have been linked to the rate of protein evolution: aligned protein length; the regional density of protein-coding bases; the density of highly conserved, vertebrate elements; and GC content (Methods V). Most of these metrics do not differ between PIPs and other genes (Fig 2 E-H), with the exception of conserved element density, which is slightly but significantly lower in PIPs (mean=8.0% vs. 8.8%; p=0.0004; Fig 2G).

Based on these results, we expanded our permutation test to find matched controls for each PIP. Control genes were considered acceptable matches if their values for each of the five significantly different metrics (Fig 2A-D; G) fell within specific ranges of the PIP value (Methods VI). This permutation procedure effectively equalized PIP and control distributions for all eight functional and evolutionary properties displayed in Fig 2A-H (S4 Fig). On average, each PIP could be matched to 32 control genes, allowing many different sets of matched controls to be generated. About 9% of PIPs were too dissimilar from other proteins to be matched, and were excluded from subsequent analysis.

Finally, one of the largest differences between PIPs and other proteins is the frequency with which they are discussed in the scientific literature (Fig 2I). The average PIP has 6.5 times more PubMed citations, and 9.1 times more scientist-contributed References Into Function (GeneRIFs), than the average mammalian protein (Methods V). This difference was too large to control for in the matched permutation test without excluding the majority of PIPs. However, we show that the citation frequency of non-PIPs has no relationship with measures of protein adaptation (p≥0.3; S5 Fig, A&B). Furthermore, non-PIPs in the top quintile of citation frequency have no more adaptation than other genes (p≥0.25; S5 Fig, C&D). This indicates that a high rate of citation for PIPs is not likely to be causally associated with their rate of adaptation.

### PIPs have experienced accelerated rates of adaptation in mammals

If malaria parasites (or similar pathogens) have indeed imposed a major selective pressure on their mammalian hosts, we would expect PIPs to exhibit unusual, adaptive patterns of amino acid substitution. In the absence of exceptional selection, however, these patterns would not be expected. After controlling for various metrics of function and genomic context (S4 Fig), we noted that PIPs have the typical ratio of non-synonymous to synonymous polymorphism in great apes (Fig 3A; mean pN/(pS+1) = 0.21 vs 0.20; p=0.40; see Enard et al., 2016). That is, PIPs do not appear more or less evolutionarily constrained than other proteins, bolstering the null expectation that they should evolve at average rates.

**Fig. 3.**
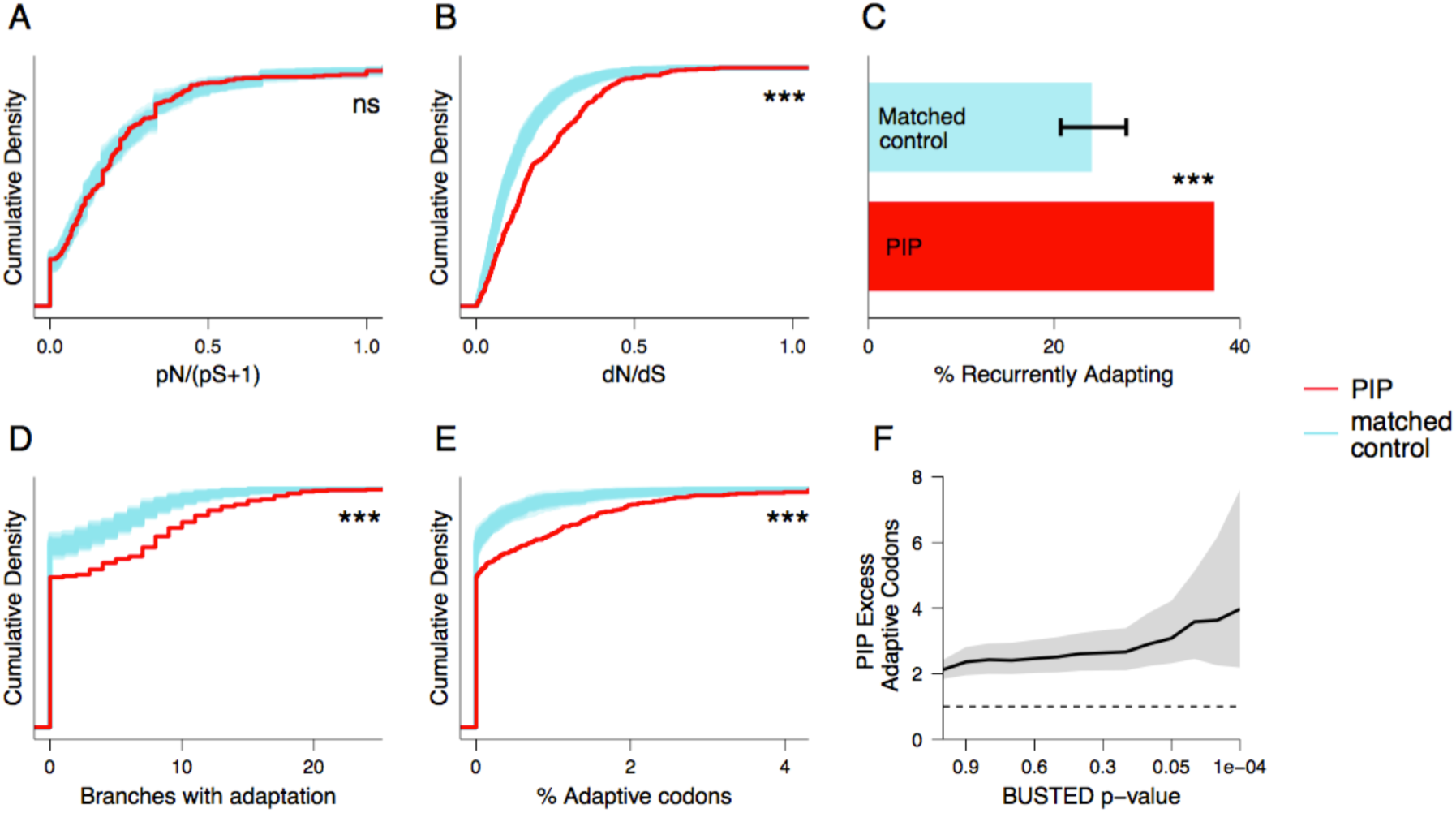
PIPs have significantly more mammalian adaptation than matched control proteins. (a) PIPs have evolved in great apes under similar levels of constraint as other proteins, as measured by extant polymorphism. (b) PIPs have 39% more non-synonymous substitutions, relative to synonymous substitutions, across 24 mammal species. (c) BUSTED detects recurrent adaptation in a greater fraction of PIPs than in sets of matched controls. (d) BS-REL tests identify PIPs as evolving adaptively on more branches in the mammalian tree. (e) BS-REL tests identify a higher proportion of codons in PIPs as evolving adaptively. (f) The excess of adaptive codons in PIPs increases (p=0.001) as the BUSTED threshold for including BS-REL estimates becomes more stringent (Methods VII). The solid line indicates the mean excess; the dashed line indicates the 1:1 expectation; gray shading indicates 95% confidence intervals. In all panels, *** = p<0.001; ns = p>0.05.

In contrast, we find that PIPs do have a significantly higher ratio of non-synonymous to synonymous substitutions across 24 mammal species (Fig 3B; mean dN/dS = 0.195 vs 0.140, p<10^−5^). This ~40% elevation in dN/dS, despite unremarkable pN/(pS+1), conservatively suggests that over a third of substitutions in PIPs may be related to positive selection imposed by malaria (or related pressures).

We refined this result using the BS-REL and BUSTED tests (Methods VII), available in the HYPHY package (Kosakovsky Pond et al., 2011; Murrell et al., 2015; Pond et al., 2005). Both tests use maximum likelihood models to estimate the proportion of codons in a protein with dN/dS > 1, consistent with adaptation. For each protein, BUSTED estimates this adaptive proportion across the entire phylogenetic tree, while BS-REL estimates it per branch

Both models find evidence of excess adaptation in PIPs. Over 37% of PIPs have BUSTED evidence (at p≤0.05) of recurrent adaptation in mammals, versus 23% of matched controls (p<10^−5^; Fig 3C). Similarly, PIPs have BS-REL evidence for adaptation on more branches of the mammalian tree (p=1.87× 10^−4^; Fig 3D), and for more codons per protein (p<10^−5^; Fig 3E). This excess is robust to the BUSTED p-value threshold used to define adaptation, and increases as the threshold becomes more stringent (Fig 3F, p=0.001). Overall, these matched tests show that PIPs have indeed experienced an accelerated rate of adaptive substitutions, consistent with malaria as an important selective pressure.

### *High rate of adaptation in PIPs known to interact only with* Plasmodium

We have shown that a large set of host proteins with strong connections to *Plasmodium* (STable 1, Fig 1 A-C) have, over deep time scales, evolved under exceptionally strong positive selection (Fig 3). Given that nearly half of PIPs are known to also interact with viruses and/or bacteria (Fig 1D), one critical question is whether *Plasmodium* is truly the source of this selection. We attempted to isolate *Plasmodium* as a selective pressure by dividing PIPs into ‘*Plasmodium*-only’ and ‘multi-pathogen’ categories, based on the available information regarding viruses and bacteria (Fig 1D; Methods IV). We find that *Plasmodium*-only PIPs have a 2.2-fold excess of adaptation compared to matched controls (p=0.008; Fig 4A, far left), when adaptation is measured as the proportion of adaptive codons per gene (Fig 3E). This suggests that *Plasmodium* may have specifically driven adaptation in a large number of mammalian proteins, apart from any pleiotropic interactions they may have with other pathogens.

**Fig. 4.**
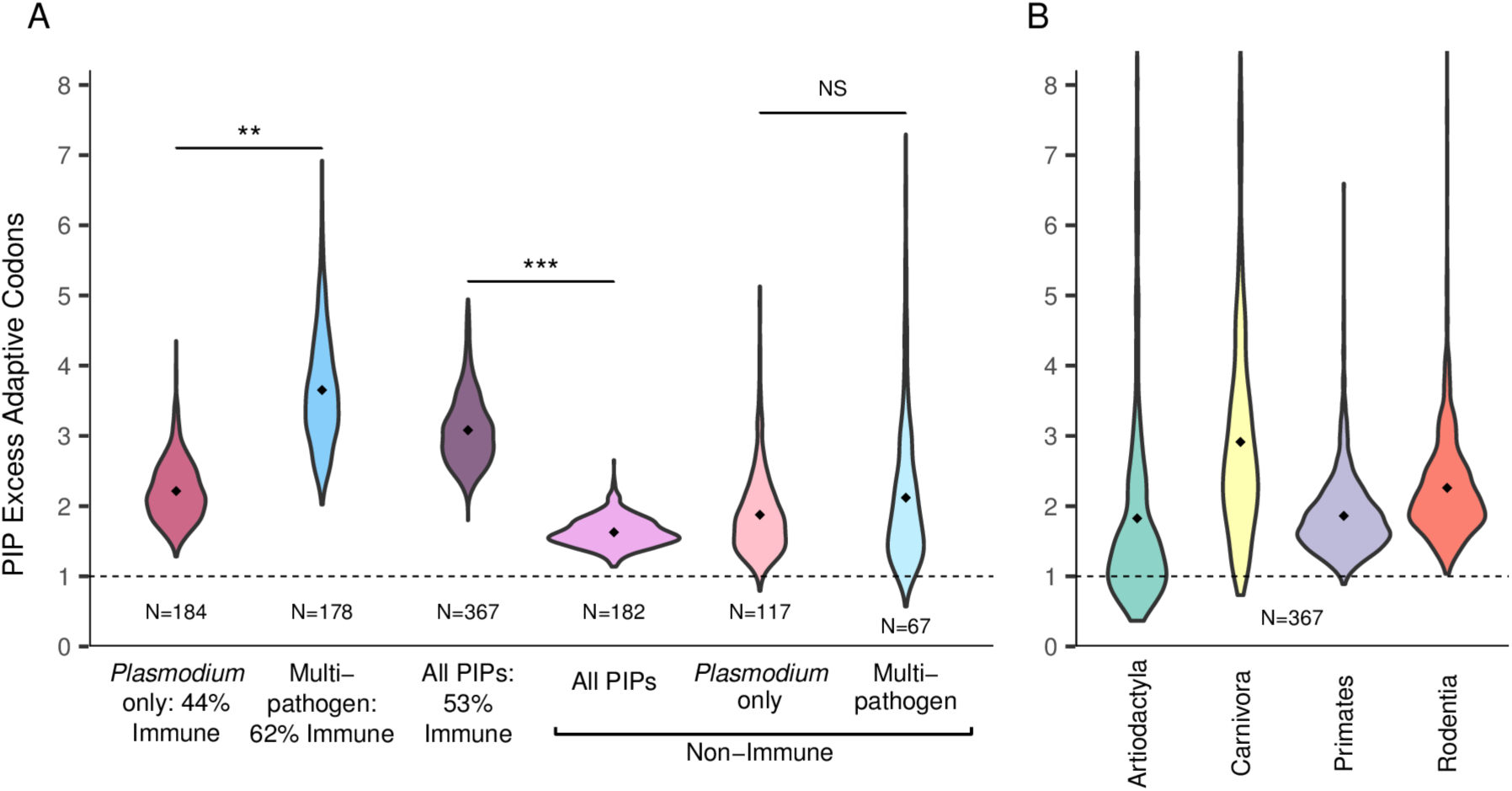
PIP adaptation is pleiotropic and pervasive. (a) Left: PIPs that interact with *Plasmodium* alone have 2.2 times more adaptation than expected. PIPs that also interact with another viral and/or bacterial pathogen have 3.7 times more adaptation than expected. Middle: Non-immune PIPs have less adaptation than all PIPs. Right: The difference between Plasmodium-only and multi-pathogen PIPs disappears when only non-immune PIPs are considered. All tested groups of PIPs exceed the 1:1 null expectation (dashed line) with p<0.05. Each violin represents the ratio between PIPs and matched controls over 1000 iterations, with the black point indicating the mean. Key: *** = p<0. 01; ** = p<0.01; NS=not significant. (b) Evidence of excess adaptation in PIPs is observed in all tested orders of the mammalian tree. The ratio is significantly higher than 1 (p<0.05) in all orders but Artiodactyla (p=0.28).

Nonetheless, multi-pathogen PIPs have 3.7 times more adaptation than matched controls—significantly higher than the excess in *Plasmodium*-only PIPs (p=0.005; Fig 4A, left). This suggests that an increased number and diversity of pathogen interactions may drive a cumulative increase in host adaptation.

Importantly, however, these multi-pathogen interactions are concentrated in immune PIPs (Fig 1D; Fig 4A). Since immune genes are well known to evolve at accelerated rates (Hurst and Smith, 1999; Nielsen et al., 2005; Bustamante et al., 2005; Voight et al., 2006; Williamson et al. 2007; Sackton et al., 2007), this immune enrichment could confound the excess of adaptation observed in multi-pathogen PIPs.

Before disentangling this issue, we first verified the correlation between immune function and adaptation (Methods VI). We find that while PIPs overall have adapted at a 3.1-fold higher rate than matched controls, non-immune PIPs have adapted at a 1.7-fold higher rate than matched, non-immune controls (Fig. 4A, middle). This difference, which is highly significant (p<0.001), reinforces that immune enrichment could confound adaptation in multi-pathogen PIPs. To isolate these two effects, we then considered only non-immune PIPs, divided into groups by their total number of pathogen interactions (Fig 4A, right; S7 Fig). In these non-immune PIPs, in contrast to all PIPs, we find that additional interactions beyond *Plasmodium* have no additional effects on adaptation.

Together, these results suggest that adaptation in immune genes is difficult to attribute to any single selective pressure. The immune system appears to be the most efficient avenue for hosts to simultaneously adapt to multiple pathogens. In contrast, host adaptation to *Plasmodium* is apparent through both immune and non-immune pathways (Fig 1D; Fig 4A). We have shown that non-immune genes evolve more slowly and have less pathogen pleiotropy (Fig 4A; Fig 1D). Thus, though *Plasmodium* has likely played a major role in immune evolution, we can be more confident that selection imposed by *Plasmodium* has specially driven adaptation in non-immune PIPs.

### PIP adaptation is related to expression in blood, liver, and lung

Malaria infections are biologically complex, and host adaptation to *Plasmodium* could occur in genes expressed in several malaria-relevant tissues (Fig 1B). We used multiple linear regression to test whether the rate of adaptation in a gene, as measured by BS-REL and BUSTED, was related to its tissue-specific expression, as measured by GTEx

For PIPs, rates of adaptation are significantly and positively related to relative expression in blood, liver, and lung, but not in other malaria-related tissues (Table 1, column 2). Overall, in a multiple linear model, PIP expression in these tissues explains 17.4% of the variance in the proportion of adaptive codons. In contrast, the tissue-specific expression of matched control genes (Methods III) explains only 4.6% of this variance in adaptation, or 3.8 times less (p<0.001). When compared to samples of control genes matched for total expression, as well as for expression in blood, liver, and lung, PIP relationships between adaptation and tissue expression are significantly stronger than expected (Table 1, column 3). This suggests that blood, liver, and lung, among all sites of PIP expression, may experience the strongest selective pressures from *Plasmodium* parasites.

**Table 1.** PIP adaptation is linked to tissue-specific expression.

| Tissue | p-value,<br>PIP adaptation ~ expression | p-value,<br>PIP vs. matched controls |
| --- | --- | --- |
| blood | $7.90 \times 10^{-3}$ | 0.012 |
| liver | $9.47 \times 10^{-16}$ | <0.001 |
| lung | $3.29 \times 10^{-4}$ | <0.001 |
| lymphocytes | 0.378 | 0.488 |
| spleen | 0.193 | 0.419 |
| s. adipose | 0.658 | 0.584 |
| v. adipose | 0.380 | 0.302 |

Relative expression in the blood, liver, and lung is more strongly related to the proportion of adaptive codons in PIPs than in other genes. 1000 sets of matched controls were matched to PIPs for total expression, as well as for relative abundance of expression in blood, liver, and lung. Abbreviations: *s* – subcutaneous; *v* – visceral.

### *PIP adaptation is not limited to* Plasmodium-*infected lineages*

A number of *Plasmodium* species infect mammalian hosts in the orders Primates and Rodentia (Carlton, Perkins, and Deitsch, 2013). In contrast, Artiodactyla and Carnivora are parasitized by other genera of Apicomplexan parasites, which also reproduce in the blood and are transmitted by insects (Clark and Jacobson, 1998). To further test the specificity of PIP adaptation, we applied the BUSTED and BS-REL models to separate protein alignments for each mammalian order (Methods VIII).

When all PIPs are considered, we find significant excesses of adaptation in rodents (p<0.001), primates (p=0.005), and carnivores (p=0.02; Fig 4B). The signal is positive, but not significant, in artiodactyls (p=0.28; Fig 4B). Artiodactyls are the most poorly-represented group in our mammalian tree (S1 Fig), and we observe that noise in these estimates is negatively correlated with the number of sampled species (R^2^=0.65; S8 Fig). The signal also becomes noisier, especially for carnivores, when the analysis is limited to *Plasmodium-only* PIPs (S9 Fig). Overall, we find no statistical evidence for the restriction of PIP adaptation to certain mammal lineages, consistent with the phenotypic similarity between *Plasmodium* and other Apicomplexan pathogens. Other ubiquitous pathogens that interact with PIPs, namely viruses and bacteria (Fig 1D), may further contribute to these mammal-wide patterns.

### *Understanding a single case of adaptation to* Plasmodium

We have shown that *Plasmodium* has driven, at least in part, an accelerated rate of adaptation in a set of 410 mammalian PIPs. In order to understand this adaptation at a more mechanistic level, we selected a single PIP for more detailed investigation.

Of the top ten PIPs with the strongest BUSTED evidence of adaptation, only one candidate—alpha-spectrin, or *SPTA1*—has been extensively characterized for molecular interactions with *Plasmodium* proteins. Alpha-spectrin is a textbook example of a major structural component of the red blood cell (RBC) membrane. In humans, several polymorphisms in this gene are known to cause deformations of the RBC, which may either be symptomless or cause deleterious anemia (reviewed in, e.g., Gallagher, 2004). The *SPTA1* protein has a well-defined domain structure, and specific interactions with *Plasmodium* proteins are known for three domains (Fig 5). Repeat 4 is the binding site for KAHRP, the major *P. falciparum* component of the adhesive ‘knobs’ that form on the surface of infected RBCs (Pei et al., 2005). Another 65-residue fragment containing EF-hand 2 has been shown to bind to PfEMP3; this interaction destabilizes the RBC skeleton, potentially allowing mature merozoites to egress from the cell (Pei et al., 2007). A central SH3 domain can also be cleaved by a promiscuous *Plasmodium* protease called plasmepsin II (Le Bonniec et al., 1999), which mainly functions in hemoglobin digestion (Francis et al., 1997). Furthermore, naturally occurring mutations in the first three *SPTA1* domains have been shown to impair the growth of *P. falciparum* in human RBCs (Schulman *et al*. 1990; Facer, 1995; Dhermy et al. 2007).

**Fig. 5.**
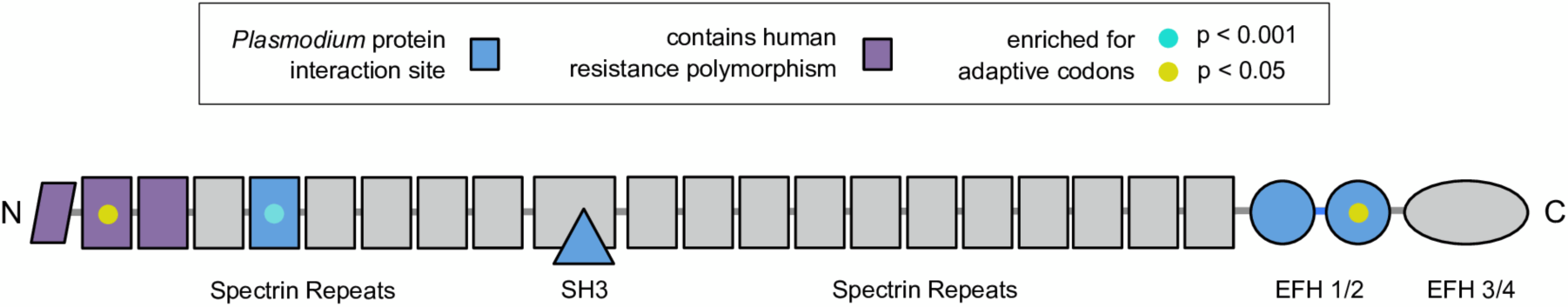
Domains of the alpha-spectrin (*SPTA1*) protein that are enriched for mammalian adaptation overlap with *Plasmodium* interaction sites. Adaptive codons were determined with MEME on an 85-species alignment of *SPTA1* coding sequence (Methods IX). *Plasmodium* interaction sites, and sites of human resistance mutations, were drawn from the literature (see text).

We wished to test whether sites of mammalian adaptation in *SPTA1* mapped to any of these *Plasmodium*-relevant domains. To identify adaptive codons with higher precision and power, we aligned *SPTA1* coding sequences from 61 additional mammal species (S5 Table) for analysis in MEME (Murrel et al., 2012; Methods IX). Of the 2,419 codons in this large mammalian alignment, we found that 63 show strong evidence of adaptation (p<0.01), and that these are distributed non-randomly throughout the protein.

Remarkably, three domains—Repeat 1, Repeat 4, and EF-hand 2—are significantly enriched for adaptive codons, after controlling for domain length and conservation (Fig 5; Methods IX). That is, all three *SPTA1* domains with strong evidence of adaptation in mammals are known to either interact specifically with *P. falciparum* proteins, or harbor human mutations that provide resistance to *P. falciparum.* This overlap is unlikely to occur by chance (p=0.015), and is robust to the p-value thresholds chosen (S6 Table). Thus, evidence from *SPTA1* suggests a meaningful and specific connection between host adaptation and the mechanics of *Plasmodium* infection.

## Discussion

In this work, we have examined decades of malaria literature to expand the collection of mammalian, *Plasmodium*-interacting proteins by over an order of magnitude (Fig 1). We show that, compared to control proteins matched for various properties (Fig 2), these 410 PIPs have adapted at exceptionally high rates in mammals (Fig 3). The highest rates of adaptation are evident in immune PIPs, especially those that share interactions with viruses and bacteria (Fig 4A). However, we show that *Plasmodium* itself (or related Apicomplexans) has likely been an important driver of this adaptation, especially for non-immune proteins (Fig 4A).

We used collections of available data on other pathogens to isolate a set of PIPs that, to the best of our knowledge, lack any ‘multi-pathogen’ interactions. These ‘*Plasmodium*-only’ PIPs, whether immune or not, have adapted at over twice the expected rate in mammals (Fig 4A). This suggests that *Plasmodium* has had an appreciable effect on PIP evolution, beyond the effect of unrelated pathogens. Still, many interactions with other pathogens likely remain unknown, making it difficult—based on this evidence alone—to dismiss their importance.

However, two other pieces of evidence support *Plasmodium* as a key selective pressure. First, mammal-wide adaptation in PIPs is strongly linked to PIP expression in human blood, liver, and lung (Table 1). *Plasmodium* parasites are well known to replicate within red blood cells (RBCs) and hepatocytes, and infected RBCs tend to sequester in the lungs, with serious consequences (e.g. Aursudkij et al., 1998). Thus, the pathophysiology of malaria is reflected in the tissues where PIPs show the strongest evidence of adaptation.

Second, in the well-studied case of alpha-spectrin, we show that domain-level interactions with *Plasmodium* perfectly explain the observed patterns of adaptive substitution (Fig 5). Besides validating the ability of codon evolution models to detect adaptation at particular residues (Methods VII), this result affirms a specific role for *Plasmodium* in mammalian evolution, beyond the immune-focused role played by pathogens in general (Fig 4A). Thus, despite the inevitability of at least some pleiotropy (Wagner and Zhang, 2011), we show that phenotypic information can be leveraged to link genetic adaptation to specific sources of selection.

Throughout this work, we showcase the utility of phenotypic information for studying evolution. We demonstrate that recent, well-funded projects like GTEx and Encode can provide, among many other uses, the raw information required for meaningful evolutionary comparisons (Fig 2; Methods). Smaller-scale projects, including most of the scientific papers contained in PubMed, also contain an impressive quantity of valuable data (Fig 1A). However, we find that heavy manual curation is still required to remove false positives from literature searches (Methods I). In the future, unique and automatic indexing of existing data will be key to understanding the evolution of complex phenotypes, and should be a major research focus, alongside the accessibility of new data.

Finally, this work provides an interesting contrast with previous studies, which have associated only a few dozen human genes with malaria resistance (Verra et al., 2009). Only a handful of these genes are backed by convincing evidence of positive selection in humans, and nearly all of these are RBC proteins (Hedrick, 2011; MalariaGEN, 2015). In contrast, our work provides a repository of hundreds of diverse human genes with phenotypic links to malaria (Fig 1; S1 Table). Why, then, do we know of so few examples of recent human adaptation to *Plasmodium*? This disconnect may depend simply on the timescale of human evolution, which is only a fraction of the ~105 million years of mammalian evolution (Murphy et al., 2007). Or, perhaps the difficulty of detecting balancing selection (Charlesworth, 2006) has obscured additional, important human variants. Future work will utilize the large set of PIPs to better understand the evolution of malaria resistance in humans.

In conclusion, we have found evidence of substantially accelerated adaptation in mammalian proteins that interact with *Plasmodium*. In the case of rapidly evolving immune proteins, *Plasmodium* appears to share responsibility with other groups of pathogens, including viruses and bacteria. We show that it can be difficult to attribute evolutionary changes to a single selective agent, given the surprising pleiotropy of host genes with regard to very different pathogenic agents. But in many cases—as well as in the case of alpha-spectrin—our approach allows us to infer that *Plasmodium*-like parasites have imposed a substantial selective pressure on mammals. We hope that our collection of 410 mammalian PIPs will continue to prove a powerful resource for exploring host interactions with *Plasmodium*.

## Methods

### I. Identification of PIPs

We queried PubMed for scientific papers containing both a gene name and the word(s) ‘malaria,’ ‘*falciparum*,’ or ‘*Plasmodium*’ in the title or abstract, as of May 21, 2015. Human gene names were drawn from the HUGO Gene Nomenclature Committee (Gray et al., 2015; <u>http://www.genenames.org/</u>) for 9,338 mammalian orthologs (Methods II). For each of the 2,249 genes that returned a hit, we manually evaluated the titles of up to 20 associated papers to assess the link between the gene and a malaria phenotype. Many acronyms used to represent genes are also used as abbreviations for techniques, locations, drugs, or other phrases. Consequently, most genes could be eliminated based on their nominal connection with papers addressing non-genetic aspects of malaria.

For papers discussing genes, we examined the abstracts for the presence and type of evidence connecting genes to malaria phenotypes. In cases where the abstract was ambiguous, we examined the full text of the paper. To limit the number of false positives, we did not include results from RNAseq or other high-throughput experiments.

### II. Generation of mammalian ortholog alignments

We used BLAT to identify homologs of 22,074 human coding sequences in 24 high-depth mammal genomes (S1 Fig). We retained orthologs which (1) had best reciprocal hits in all 24 mammal species, (2) lacked any in-frame stop codons, (3) were at least 30% of the length of the human sequence, and (4) had clearly conserved synteny in at least 18 non-human species. Coding sequences for the resulting 9,338 proteins were aligned with PRANK, and any codon present in fewer than eight species was excluded from analysis. Additional details are available in Enard et al. (2016).

### III. Tissue Expression Analyses

Expression data for 53 human tissues were downloaded from the GTEx portal (<u>http://www.gtexportal.org/home/</u>) on October 18, 2015. For tissue-specific analyses (Fig 1B, Table 1), we converted RPKM values to relative abundance (RA) values for each tissue. RA is simply the proportion of each gene’s total RPKM found in each tissue. For matching controls, we summed RPKM values over all tissues to yield total expression.

Because PIPs have substantially higher total expression than other proteins (Fig. 2A), for each PIP, we identified matched control proteins with +/−20% of the total RPKM expression of that PIP. For each of 10,000 iterations, we randomly selected one matched control out of the set of potential matches for each PIP, producing PIP and control sets of equal size and with indistinguishable distributions of total expression (S4 Fig). We compared the mean value from each matched control set to the mean value for PIPs, and determined p-values empirically, as the fraction of permutations with control mean ≥ PIP mean. Tissues with significantly different PIP expression were determined after applying the Bonferroni correction for 53 tissue tests (S2 Fig).

We also correlated tissue-specific expression in malaria tissues with the proportion of adaptive codons per gene (Table 1), using multiple linear regression in R. To generate matched controls for this analysis, we matched control genes to PIPs based on their RA in blood, liver, and lung, as well as total expression.

### IV. Collection of VIPs and BIPs

Virus-interacting proteins (VIPs) were manually curated by Enard *et al.* (2016), in the same manner as PIPs. To our knowledge, no similar collection of high-quality interactions is available for other pathogens. Therefore, we queried the EBI IntAct database (<u>http://www.ebi.ac.uk/intact/</u>) for protein interactions between Kingdom Bacteria (taxid:2) or Phylum Apicomplexa (taxid:5794) and humans (taxid:9606). This approach, while much faster than manual curation, is less ideal for two reasons: (1) many interactions are not included in the database (e.g., only 17 human-*Plasmodium* interactions are included in IntAct), and (2) many of the included interactions are based on high-throughput assays, including yeast two-hybrid experiments, which suffer from both false negatives and false positives (Brückner et al 2009). Consequently, we do not perform rigorous analysis for bacterial-interacting proteins (BIPs), as has been done for PIPs and VIPs (Enard et al., 2016). Rather, we use the IntAct BIPs only to classify PIPs as ‘*Plasmodium*-only’ or ‘multi-pathogen.’

### V. Collection of other protein metrics

GO annotations were downloaded in October, 2015 from the Gene Ontology website (Ashburner et al., 2000; <u>http://geneontology.org/</u>)

Regions of DNaseI hypersensitivity, combined from 95 cell types, were obtained from the databases of the ENCODE Project Consortium (2012; <u>https://www.encodeproject.org/</u>). We calculated the density of DNaseI hypersensitivity regions in 50 Kb windows centered on each ortholog.

Protein expression levels were obtained from the Human Proteome Map (Kim et al., 2014; <u>http://www.humanproteomemap.org/</u>), which used high resolution and high accuracy Fourier transform mass spectrometry experiments. We summed spectral values over 30 tissues and cell types and took the log of these total values. The log number of interacting partners for each human protein was obtained from the Biogrid Database (Stark et al., 2011; <u>http://thebiogrid.org/</u>), curated by Luisi et al., 2015.

Genomic elements conserved in 46 vertebrate species, derived from PhastCons (Siepel et al., 2005), were downloaded from the UCSC genome browser (<u>http://hgdownload.cse.ucsc.edu/goldenPath/hg19/phastCons46way/</u>). We calculated conserved element density within 50 kb windows centered on each gene in the human reference. Coding density was calculated from coding nucleotides in the same 50 Kb windows. The length and GC content of each protein was derived from the mammalian alignment (Methods II).

We assessed citation frequency of each gene in two ways. First, we counted the citations linked to each gene on its PubMed Gene page (<u>http://www.ncbi.nlm.nih.gov/gene</u>). Second, we downloaded the Gene References into Function, or GeneRIFs, contributed to PubMed by scientists (<u>ftp://ftp.ncbi.nih.gov/gene/GeneRIF/</u>). These measures were highly correlated (not shown), and only citations are reported.

### VI. Matching PIPs to control proteins

Each PIP was matched to a set of control proteins based on similarity in five metrics: mRNA expression, protein expression, protein-protein interactions, DNaseI density, and conserved element density (Fig. 2A-D; G). We allowed a control protein to be considered a PIP ‘match’ if each of its five values fell within a given range, based on the PIP values. For example, margins of min=0.1 and max=0.2 for mRNA expression means that, for a control protein to be matched to a PIP, the mRNA expression of the control must fall between 90-120% of the mRNA expression of the PIP. We wished to maximize the number of matched controls per PIP while creating control sets that were statistically indistinguishable from PIPs for all five metrics (e.g. S4 Fig). To achieve this balance, we iteratively chose the maximum margins that yielded average p-values, over 100 permutations, of ≥ 0.1 for each metric. Once appropriate margins were found, we obtained matched control sets of equal size to the PIP set by randomly sampling one matched control protein for each PIP. For each permutation test, 10,000 sets of matched controls were sampled.

Margins for the main permutation test (Fig 3) are given in S4 Table. For subsets of PIPs (i.e., Fig 4A, S7 Fig), the margins were altered to generate well-matched controls in every case. For Table 1, only the stated expression values and pN/(pS+1) were checked for matching, to avoid excluding too many PIPs. Because we chose sets of matched controls based on the distribution of PIP values included in each test, whether any given PIP was matched depended on the other PIPs in the test (e.g., one extreme PIP may or may not be balanced out by another). Therefore, the sum of matched PIPs across categories differs slightly from the total (Fig 4A).

The pool of immune controls is relatively small (998 genes), compared to the pool of non-immune controls with other GO annotations (7,594 genes)(S2 Table). This made it difficult to match immune PIPs to immune controls, without discarding many immune PIPs. Consequently, to test hypotheses of accelerated immune adaptation, we focused on comparing all PIPs to all controls and non-immune PIPs to non-immune controls (Fig 4A). For these and other tests in Figure 4, 1000 sets of matched controls were sampled for each violin.

### VII. Estimating adaptation with models of codon evolution

We used the codeml model m8 from the PAML package (Yang, 2007) to estimate dN/dS for each ortholog (Fig 3B). However, branch-site tests in PAML rely on assumptions that may be violated in the case of recurrent adaptation to a pervasive selective pressure (see Enard et al. 2016). Consequently, we chose to implement the maximum-likelihood branch-site tests in the better-performing HYPHY package (Kosakovsky Pond et al., 2011). We used the BUSTED algorithm (Murrell et al., 2015) to detect recurrent selection across the entire tree for each gene, and BS-REL to estimate the proportion of positively selected codons in each gene on each branch. Both of these algorithms rely on the same underlying codon model; details of the model are described in Kosakovsky Pond et al. (2011), Murrell et al. (2015), and reviewed in Enard et al. (2016). Unless otherwise specified (i.e., Fig 3F), codons identified by BS-REL were ‘counted’ as adaptive if the BUSTED p-value for that gene was ≤0.05. For each evolutionary statistic (i.e. adaptive codons, adaptive branches, dN/dS, pN/pS), empirical p-values were derived by comparing the average value of the PIPs to the average values of 100,000 permutations of matched controls.

### VIII. Order-specific analyses

We split the mammal-wide alignments for each gene into four non-overlapping alignments, corresponding to the following clades: **primates** (human, chimp, gorilla, orangutan, gibbon, macaque, baboon, marmoset, bushbaby), **rodents** (mouse, rat, guinea pig, squirrel, rabbit), **carnivores** (panda, ferret, dog, cat), and **artiodactyls** (sheep, cow, pig) (see S1). We excluded microbat, elephant, and horse, as these species are not closely related to any of the four major groups (Murphy et al., 2007; S1 Fig). However, we included rabbit with rodents, because they are more closely related. We ran BUSTED on each alignment to yield a p-value of clade-specific adaptation for each gene.

PIPs were matched to controls as described above (Methods V). However, rather than counting BS-REL adaptive codons in all branches if the tree-wide BUSTED p≤0.05, we (1) kept each clade codon count separate, (2) counted codons only on branches within a clade, and (3) counted codons only if the clade-specific BUSTED p≤0.05. The ratio of adaptive codons in PIPs versus controls was then calculated as before, by taking 1,000 random samples of matched controls.

### IX. Alpha-spectrin

Alpha-spectrin homologs were initially identified in 88 mammal species using NCBI Gene (<u>http://www.ncbi.nlm.nih.gov/gene/?Term=ortholog_gene_6708</u>).

The sequence of the longest mRNA transcript for each species was downloaded using E-Utilities, and each transcript was trimmed to the longest ORF using TransDecoder (Haas et al., 2013; <u>http://transdecoder.github.io/</u>). Coding sequences with <50% of the human CDS length were removed. The remaining 85 coding sequences were aligned with PRANK (Löytynoja and Goldman, 2008) using default settings (S5 Table). The alignment was manually inspected and corrected using JalView (Waterhouse et al., 2009).

A phylogenetic tree for the 85 species was obtained, using NCBI Taxonomy, from phyloT (<u>http://phylot.biobyte.de/</u>). This tree, along with the corrected alignment, was inputted into HyPhy to run MEME (Murrell et al., 2012), which yielded a p-value of adaptation for each codon. We used the domain designations from SMART (Schultz et al., 1998; <u>http://smart.embl-heidelberg.de/</u>) to assign 92.2% of *SPTA1* codons to one of 25 domains (S6 Table). Then, for each domain, we calculated an ‘adaptation score’ as:

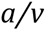

where *a* measures adaptation (the proportion of codons within the domain with MEME p≤0.01*) and *v* measures variability (the proportion of codons within the domain that vary among species, i.e., are not 100% conserved). This score also controls for domain length, as it uses the proportion of codons within the domain. To calculate the significance of each domain’s adaptation score (i.e., to ask, is it higher than expected?), we randomly permuted codons among domains 10,000 times.

*We also tested MEME p-value cutoffs of 0.1, 0.5, 0.005, and 0.001 for defining *a*; these results are available in S6 Table. The results for p≤0.01, which are reported in the main text, are representative across these cutoffs.

## Data Access

All data used in this work are publicly available (Methods I-V). The collection of PIPs is available in S1 Table.

## Acknowledgements

We wish to thank Kerry Geiler-Samerotte for her thoughtful comments on the manuscript, along with the rest of the Petrov lab. ERE thanks Jane Carlton for abbreviation advice; Jamie Blundell and Anisa Noorassa for figure advice; and Daniel Friedman, for conceding that ‘protein’ can mean ‘gene.’

## Author Contributions

E.R.E curated the PIPs. D.E., E.R.E., and N.T. collected other data. E.R.E. and D.E. performed the analyses, with design input from D.A.P. and S.V. E.R.E. and D.A.P. wrote the paper, with contributions from all other authors.

This work was supported by NIH grants R01GM089926 and R01GM097415 and NSF grant R35GM118165-01 to DAP, and an NSF Graduate Research Fellowship to ERE (DGE-1247312).

